# Autotrophic biofilms sustained by deeply-sourced groundwater host diverse CPR bacteria implicated in sulfur and hydrogen metabolism

**DOI:** 10.1101/2022.11.17.516901

**Authors:** Luis E. Valentin Alvarado, Sirine C. Fakra, Alexander J. Probst, Jonathan R. Giska, Alexander L. Jaffe, Luke M. Oltrogge, Jacob West-Roberts, Joel Rowland, Michael Manga, David F. Savage, Chris Greening, Brett J. Baker, Jillian F. Banfield

## Abstract

**Background:** Candidate Phyla Radiation (CPR) bacteria are commonly detected yet enigmatic members of diverse microbial communities. Their host associations, metabolic capabilities, and potential roles in biogeochemical cycles remain under-explored. We studied chemoautotrophically-based biofilms that host diverse CPR bacteria and grow in sulfide-rich springs using bulk geochemical analysis, genome-resolved metagenomics and scanning transmission x-ray microscopy (STXM) at room temperature and 87° K.

**Results:** CPR-affiliated Gracilibacteria, Absconditabacteria, Saccharibacteria, Peregrinibacteria, Berkelbacteria, Microgenomates, and Parcubacteria are members of two biofilm communities dominated by chemolithotrophic sulfur-oxidizing bacteria including *Thiothrix* or *Beggiatoa*. STXM imaging revealed ultra-small cells along the surfaces of filamentous bacteria that we interpret are CPR bacterial episymbionts. STXM and NEXAFS spectroscopy at carbon K and sulfur L_2,3_ edges show protein-encapsulated elemental sulfur spherical granules associated with filamentous bacteria, indicating that they are sulfur-oxidizers, likely *Thiothrix*. Berkelbacteria and Moranbacteria in the same biofilm sample are predicted to have a novel electron bifurcating group 3b [NiFe]-hydrogenase, putatively a sulfhydrogenase, potentially linked to sulfur metabolism via redox cofactors. This complex could potentially underpin a symbiosis involving Berkelbacteria and/or Moranbacteria and filamentous sulfur-oxidizing bacteria such as *Thiothrix* that is based on cryptic sulfur cycling. One Doudnabacteria genome encodes adjacent sulfur dioxygenase and rhodanese genes that may convert thiosulfate to sulfite. We find similar conserved genomic architecture associated with CPR bacteria from other sulfur-rich subsurface ecosystems.

**Conclusions:** Our combined metagenomic, geochemical, spectromicroscopic and structural bioinformatics analyses link some CPR bacteria to sulfur-oxidizing Proteobacteria, likely *Thiothrix*, and indicate roles for CPR bacteria in sulfur and hydrogen cycling.

## Background

Sulfur is the fifth most abundant element on earth and the sulfur cycle is a key component of Earth’s interlinked biogeochemical cycles[1,2]. In natural ecosystems, sulfur exists in several oxidation states, -2, 0, +2, +4 and +6 being the most common, in the forms of polysulfide (HS_x_ or S _x_^2-^ ; -2,0), thiosulfate (S_2_ O_3_^2-^ ; -1,+5), tetrathionate (S_4_ O ^62-^ ; -2,+6), sulfite (SO_3_^2-^ ; +4) and sulfate (SO_4_^2-^ ; +6). Microbes play an important role in sulfur cycling in aqueous and soil environments. H_2_ S is also a toxic compound that must be maintained at low levels for the sustained growth of microbial consortia, thus microbial sulfide oxidation is beneficial at the community level.

Sulfide (S^2-^) is common in natural springs and can serve as a source of energy and reducing power for chemolithoautotrophic microorganisms. Chemolithoautotrophic microbial communities with members that carry out the oxidation, reduction and disproportionation of sulfur compounds are found in environments such as hydrothermal vents[3,4], water column oxic/anoxic interfaces[5–7], terrestrial caves[8–10], groundwater[11,12] and activated sludge[13]. Sulfur-based chemoautotrophic cave mats are dominated by filamentous *Campylobacterota* in environments with high S^2-^/O_2_ (>150) ratios, whereas *Gammaproteobacteria* (*Beggiatoales* and *Thiothrixales*) are prevalent at lower S^2-^/O_2_ (<75) ratios[9]. *Beggiatoaceae* and *Thiotrichaceae* that have been cultivated have been shown to use hydrogen sulfide either mixotrophically or heterotrophically [14–17]. *Beggiatoa spp*. are gliding filamentous bacteria that form S^0^ spherical granules that they may oxidize to sulfate when H_2_ S supply becomes limited [18]. *Thiotrix spp*. are gliding bacteria that can grow as long filaments (cells in a microtubular sheath) and are known to accumulate S^0^ spherical granules when in the presence of reduced sulfur[13,19] and organics (energy and carbon source) [14]. Prior work[20–24] indicate that sulfur-oxidizing bacteria support communities by providing resources such as fixed carbon and nitrogen.

To date, most studies of sulfur-based chemoautotrophic ecosystems have investigated the roles of the relatively most abundant organisms. However, it is well understood that microbial biofilms are structured as networks of interacting organisms, some of which are fundamentally dependent on other community members. Of particular interest are Candidate Phyla Radiation (CPR) bacteria (also known as Patescibacteria) [25–28] that can form symbioses with host organisms [29–31]. Prior surveys have documented CPR bacteria in sulfur-based communities [25,32,33], yet the nature of CPR-host relationships and the roles of CPR in sulfur-based communities remain under-explored.

Here, we studied chemoautotrophic microbial communities sustained by sulfur metabolism in two mineral springs MS4 and MS11[34] at Alum Rock Park, CA, USA, where sulfide-rich groundwater discharges along the Hayward fault. We profiled oxygen isotopes, temperature, water composition and spring discharge rates to constrain the sources of water and further combined genome-resolved metagenomics with electron microscopy and X-ray spectromicroscopy to investigate metabolic capacities, interdependencies, and structure of the microbial biofilm community at these two springs. Synchrotron-based spectromicroscopy evidenced the close association between ultra-small cells, inferred to be CPR bacteria, and sulfur-oxidizing bacteria that underpin this chemoautotrophic ecosystem. We predict the contributions of the major community members to carbon, nitrogen, hydrogen and sulfur cycling and investigate the potential roles of the abundant and diverse CPR bacteria in these consortia.

## Materials and Methods

### Site Description and Microbial biomass collection

The spring system is located along Penitencia Creek in Alum Rock Park, San Jose, CA (37°23’57.7”N, 121°47’48.8”W) **(Fig. 1A)**. The two sample sites, Mineral Springs 4 and 11 (MS4 and MS11) are located on opposite sides of the creek approximately 250 m from one another (**Fig. 1B-C**). Samples for geochemical analyses and 16S rRNA gene sequencing were taken in May 2005, during the dry season, and were filtered on-site using sterile 0.2 µm filters. Biofilm samples for scanning electron microscopy were collected from both sites using sterile pipettes. Solutions were acidified with 3% nitric acid for cation analyses. Samples were transported back to the laboratory on ice. Biofilm samples for metagenomic sequencing were collected on November 1, 2012 and July 2, 2019 and July 24, 2020. Planktonic samples were collected June 10, 2015 and July 24, 2020. Two sets of planktonic samples were taken by sequentially filtering 379 L and 208 L of water, respectively, from the MS4 spring onto 0.65 μm and 0.1 µm large volume filters (Gravertech 5 inch ZTEC-G filter). Filters were frozen on dry ice at the site and stored at -80°C for genome-resolved metagenomic analyses. For synchrotron measurements (STXM and X-ray microprobe), thin white streamers were collected in June 2015 with sterile tweezers at both sites and transported in falcon tubes on ice. Samples were then thawed and immediately deposited either onto a Si_3_ N_4_ window (TEM windows) or a Cu TEM grid (300 mesh, Ted Pella). Samples were then plunged in liquid nitrogen for cryogenic measurements, gas ethane (used for flash-freezing) was not available at the time of sampling. For all synchrotron-based measurements, samples were not rinsed or spinned so as to preserve the structural integrity of the filaments and preserve the CPR bacteria-bacteria-filaments spatial relationships.

**Figure 1.**
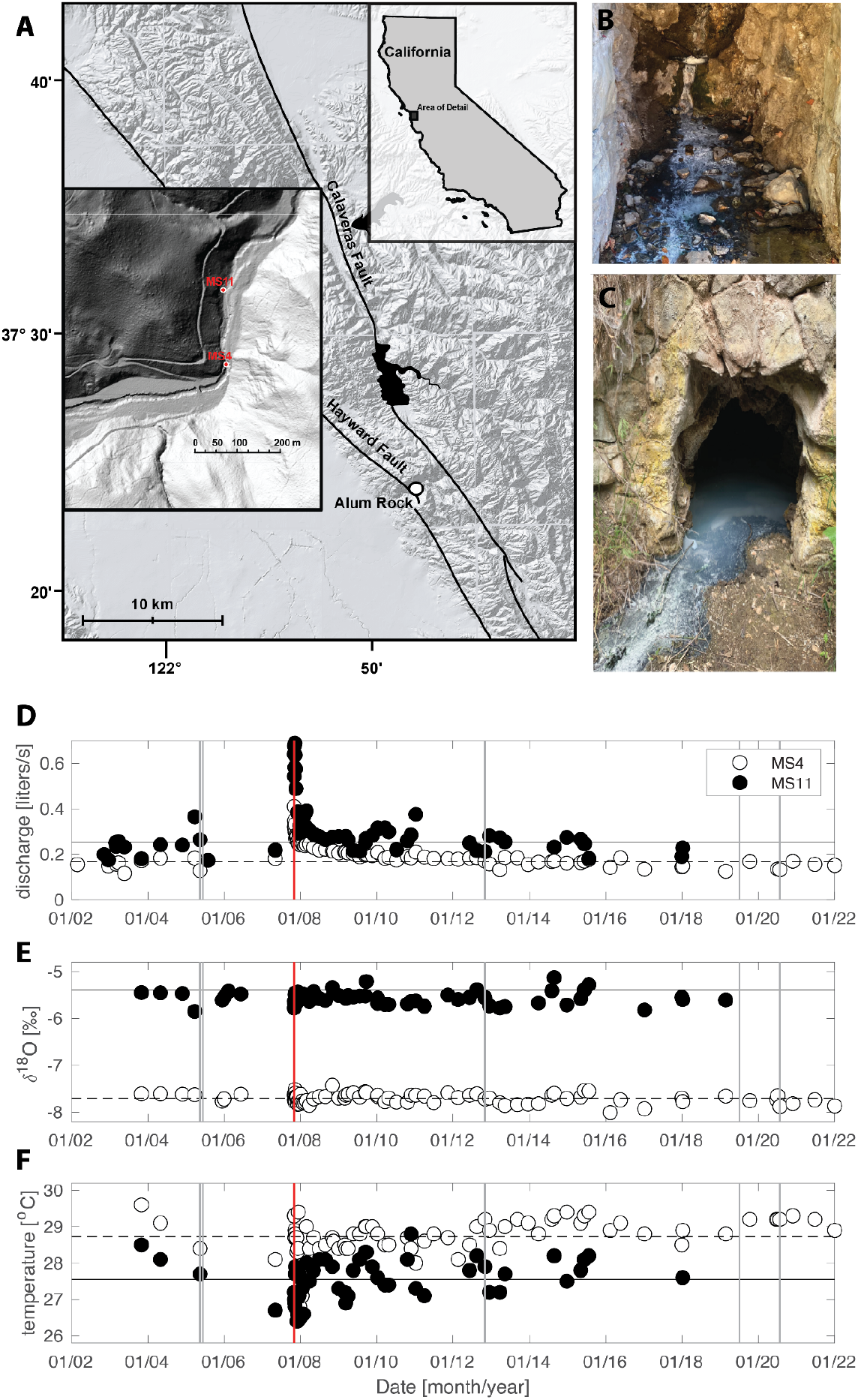
A) Shaded relief map showing the location of Alum Rock springs, CA, USA. Insets show the location of Alum Rock and of the MS4 and MS11 springs. Photographs of B) MS4 and C) MS11 biofilms. Thin white streamers (5-10 cm) are mostly found attached to the surfaces of rocks. Hydrogeological properties D) Discharge E) δ^18^Ο, and F) Temperature are steady over periods greater than a decade, except following large regional earthquakes. A discharge increase in late 2007 followed a magnitude 5.6 earthquake with an epicenter 4 km from the springs (vertical red line), neither δ^18^Ο nor the temperature changed indicating that fluid sources did not change. The horizontal lines show averages of plotted quantities over the entire sampling period, except discharge for which the average excludes the first two years after the earthquake. Vertical grey lines show dates of biofilm and planktonic sampling.

### Geochemical Analysis

Water discharge (volume/time) was measured by diverting water into either a bucket or graduated cylinder to measure volume, and time was recorded with a stopwatch. Temperature was measured with a type K thermocouple until February 2008 and thereafter with a thermistor. Accuracy is 0.2 °C and 0.1 °C, respectively. Water for O and H isotope measurements was collected in 250 mL Nalgene bottles. Discharge and temperature were not measured if outflow channels from the springs backed up to create pools of water. Cation analysis was performed on a PerkinElmer 5300 DV optical emission ICP with autosampler. Anion analysis was performed on-site using a HACH DR2010 spectrophotometer with protocols provided by the manufacturer. O and H isotopes were measured with a GV IsoPrime gas source mass spectrometer, with analytical precision of approximately 0.1 and 1 permil, respectively.

### Scanning Electron Microscopy

Scanning electron microscopy samples were fixed for two hours in a 2% glutaraldehyde solution (in 0.1 M sodium cacodylate buffer) according to a standard protocol, then vacuum aspirated onto0.22 µm polycarbonate filters (Osmonics, poretics, 47 mm, Catalog number K02CP04700), and rinsed three times in 0.1 M sodium cacodylate buffer. The samples were then dehydrated in successive ethanol baths of increasing concentration and finally dried using a Tousimis AutoSamdri 815 Critical Point Dryer for approximately one hour. Specimens were mounted on gold stubs and sputter coated with a gold/palladium mix. Imaging was performed on a Hitachi S-5000 scanning electron microscope at 10 keV at UC Berkeley.

### Scanning Transmission X-ray Microscopy (STXM)

STXM and near edge x-ray absorption fine structure (NEXAFS) spectroscopy measurements were performed on the soft X-ray undulator beamline 11.0.2[35] of the Advanced Light Source (ALS), Berkeley, CA, USA. Data were recorded with the storage ring operating in top-off mode at 500 mA, 1.9 GeV. Samples were thawed right before STXM-NEXAFS measurements at ambient temperature under He at pressure <1 atm. A Fresnel zone plate lens (40 nm outer zones) was used to focus a monochromatic soft X-ray beam onto the sample. The sample was raster-scanned in 2D through the fixed beam and transmitted photons were detected with a phosphor scintillator-photomultiplier assembly; incident photon counts were kept below 10 MHz. The imaging contrast relies on the excitation of core electrons by X-ray absorption [36–38]. STXM images recorded at energies just below and at the elemental absorption edge (S L_3_ and C K) were converted into optical density (OD) images where the OD for a given energy can be expressed from the Beer-Lambert law, for a given X-ray energy, as OD= -ln(I/I_0_)= µ *ρ* t, where I, I_0_, µ, *ρ* and t are the transmitted intensity through the sample, incident intensity, mass absorption coefficient, density and sample thickness, respectively. Protein, carbon and elemental sulfur maps were obtained by taking the difference of OD images at 280 and 288.2 eV, at 280 and 305 eV, and at 162 and 163.9 eV respectively. Image sequences (‘stacks’) recorded at energies spanning the S L_2,3_ -edges (160-180 eV) with steps of 0.3 eV around the L_3_ -edge, and C K-edge (280-305 eV) with steps of 0.12 eV around the K-edge were used to obtain NEXAFS spectra from specific regions. S 2p NEXAFS spectral features are affected by spin-orbit splitting and molecular field, and provide information on the oxidation state of sulfur.

Additionally, STXM-NEXAFS measurements at 87° K were performed on frozen-hydrated samples so as to preserve sample chemical and structural integrity[39] and minimize beam-induced radiation damage. These samples were cryo-transferred through a specimen chamber (<100 mTorr) into an LN_2_ -cooled stage (87°K) inside the STXM operated with a scanning Fresnel zone plate lens (60 nm outer zones), under vacuum (10^−6^ torr). With this setup, the sample is not rastered-scanned so as to minimize sample vibrations, instead the zone plate is scanned in 2D. Note that sulfur L_2,3_ -edges could not be accessed in this configuration due to geometrical constraints.

At least two different sample regions were analyzed at each elemental edge and beam-induced radiation damage was carefully checked. The theoretical spectral and spatial resolutions during measurements were +/-100 meV ; 40 nm and 60 nm respectively. The photon energy was calibrated at the C K-edge using the Rydberg transition of gaseous CO_2_ 292.74 eV (C → 1s 3s (v = 0)). Sulfur spectra were calibrated using the S 2p_3/2_ edge of elemental sulfur set at 163.9 eV. An elemental sulfur standard spectrum was kindly provided by Geraldine Sarret (University Grenoble Alpes, France). All data was processed with the aXis2000 software version 06 Jul 2021 (http://unicorn.mcmaster.ca/aXis2000.html).

### X-ray fluorescence microprobe (XFM)

Synchrotron XFM measurements were performed in cryogenic conditions (95°K) at ALS XFM beamline 10.3.2[40], with the storage ring operating in top-off mode at 500 mA, 1.9 GeV. Micro-focused X-ray fluorescence (µXRF) elemental mapping was performed on LN_2_ -frozen hydrated samples oriented at 45° to the incident X-ray beam, samples were cryo-transferred into a LN_2_ - cooled apparatus following procedures described elsewhere[41]. All data were recorded using a single-element XR-100 silicon drift detector (Amptek, Be window).

XRF maps were recorded at 4138 eV (100 eV above the Ca K-edge) using a beam spot size of 3 µm × 4 µm, 2 × 2 µm pixel size and 70 ms dwell time/pixel. Micro-XRF spectra were recorded simultaneously on each pixel of the maps. All maps were then deadtime-corrected and decontaminated using custom LabVIEW 2018 (National Instruments, Austin, TX, USA) software available at the beamline. Maps were then processed using a custom Matlab R2020b program (MathWorks, Natick, MA, USA) available at the beamline.

### DNA extraction and metagenomic sequencing

Approximately 200 µl of biofilm was extracted using MoBio Powersoil DNA extraction kit (MoBio Laboratories, Inc., CA, USA) according to the manufacturer’s protocol, with the bead-beating time reduced to less than one minute. This DNA extract was then gel purified and quantified using a low-mass ladder (Promega). PCR was performed on ∼50 ng of DNA in a reaction mixture containing 1X Takara ExTaq PCR buffer, 2 mM MgCl2, 50 µg of non-acetylated BSA, 200 µM dNTPs, 12.5 ng of universal bacterial 16S rRNA gene primers (27F and 1492R), 1.5 U ExTaq polymerase (Takara, Madison, Wisc.), and made to a volume of 50 µl with sterile milliQ water. Reactions were optimized for annealing temperature over the range of 48-60°C for 25 cycles and the most intense single bands were gel purified.

Total genomic DNA for metagenomic sequencing (150 bp or 250 bp reads) for both biofilm and planktonic samples (20% of each filter) was extracted using MoBio PowerMax Soil DNA extraction kit. Cells were extracted from 20% of each filter by adding 15 ml of lysis buffer and vortexing for 10 minutes. Lysis of cells was modified by heating to 65°C for 30 minutes and 1 min of bead beating. DNA was eluted in milliQ water and ethanol precipitation was performed (70% EtOH, 3 M sodium acetate, incubation for 24 hours at 4 °C).

### Illumina sequencing, assembly, binning and sequence curation

Shotgun genomic reads were assembled using IDBA-UD [42].Draft genomes consisting of scaffolds ≥ 1 kbp in length were binned based on a combination of GC content, coverage, single copy gene content, phylogenetic profile and patterns of organism abundance over samples. The phylogenetic profile was established using a database of isolate as well as metagenomics-derived sequences. In some cases, scaffold sequences from groups of bins were used to construct emergent self-organizing maps in which the structure was established using tetranucleotide composition (tetra-ESOMs). For scaffolds > 6 kb, scaffolds were subdivided into 3 kb segments and treated separately in the ESOM analysis. In cases where the majority of segments from the same scaffold did not group together in the ESOM, the scaffolds were evaluated manually (based on gene content and other information) to resolve their placement or assign them to unbinned. The scaffold set defined based on ESOM analysis was then used to generate a draft genome bin that was again checked for consistent binning signals (as above). As ESOMs only used scaffolds >3 kb in length, scaffolds from the original bins were added if they had a tightly defined GC, coverage and the expected phylogenetic profile. CheckM [43] was used for estimation of genome completeness, strain heterogeneity and contamination. Curated genomes with less than 5 duplicated single-copy genes (some of which occur because genes are split at scaffold ends) and with ≥ 95% of the expected single copy marker gene set used for completeness estimation (50 for CPR, 51 for other bacteria) were classified as near-complete, ≥ 70% and < 90% complete as drafts and those < 70% complete as partial. Genomes with >5 duplicated single-copy genes were classified as partial, regardless of other indicators of bin completeness. Candidate phage contigs were identified based on their lack of consistent phylogenetic profile and the presence of proteins with homology to those of known phages. Those with similar characteristics, and typical plasmid genes, but lacking typical phage structural genes were labeled as plasmids. Manually curated phages were classified using Virsorter2[44]. Other viral sequences were profiled using Virsorter2, evaluated by checkV [45] and annotated using DRAMv [46] with default parameters.

### Phylogenetic analyses

The concatenated ribosomal protein tree was generated using 16 syntenic genes that have been shown to undergo limited lateral gene transfer (rpL2, 3, 4, 5, 6, 14, 15, 16, 18, 22, 24 and rpS3, 8, 10, 17, 19) [47]. We obtained branch support with the ultrafast bootstrap [48] implemented in iQ-TREE v1.6.12 [49] with the following parameters: -bb 1000 -m LG+F+G4. Trees were visualized using iTOL v6.3.2 [50]. Amino acid alignments of the individual ribosomal proteins were generated using MAFFT v7.304 [51] and trimmed using trimAL [52] with the following setting: -gt 0.1.

To verify the presence of biogeochemically-relevant genes, phylogenetic trees were constructed. We used markers for sulfur (DsrAB, Pdo), carbon metabolism (RuBisCO) and energy conservation ([NiFe]-hydrogenases). Sequences were obtained using GOOSOS and aligned using MAFFT v7.304. The phylogeny for DsrAB was generated using FastTree 2.1.11 SSE3 [53]. All other phylogenies were generated using iQ-TREE v.1.6.12 using the ultrafast bootstrap and parameters specified previously.

Hydrogenase sequences from Alum Rock genomes were obtained using HMMs from [54]. Phylogenetic classification was performed using reference sequences obtained from [54] and using HydDB [55]. Verification of hydrogenase loci was performed via inspection of nearby genes and the presence of required hydrogenase accessory genes. Genome context diagrams were generated using Clinker[56].

### Metagenomics metabolic pathways analysis

Preliminary functional annotations were established and collections of metabolic capacities in genome bins were overviewed using ggKbase tools [57]. In addition, metabolic profiling was done by mapping ORFs to KEGG ortholog groups (KOs) using an HMM database that was compiled as previously described[58]. This HMM database was used to scan the metagenomic bins, and ORFs were assigned the KO of the best-scoring HMM, providing it was above the noise threshold. In addition, we profiled metabolic capacities with KEGG functional annotation using METABOLIC[59].

### Protein structure prediction

Protein structures were predicted for the putative complexes of the nitrate reductase (Nrx), dioxygenase/rhodonase, and group 3b [NiFe]-hydrogenase using AlphaFold2 in multimer mode. In all cases, the average per residue confidence scores (pLDDT) exceeded 90, a level that is empirically shown to produce highly accurate local structural models. The best-scoring models were aligned to related protein complexes in PyMol. Group 3b [NiFe]-hydrogenase complexes were predicted using AlphaFold2 in multimer mode for the HyhL (hydrogenase large subunit), HyhS (hydrogenase small subunit), HyhG (diaphorase catalytic subunit) and HyhB (diaphorase electron transfer subunit)[60,61].

## Results

### Groundwater of mixed origin hosts biofilms dominated by filamentous bacteria

We measured the flow rate, pH, and concentrations of ionic species (**Supplementary Table S1**) in the MS4 and MS11 groundwater. The MS11 spring has higher flow rate, ionic strength, alkalinity, and sulfide levels than the MS4 spring. H and O stable isotope compositions of the waters, combined with salinity measurements, indicate that spring waters are mixtures of meteoric input and pore waters from the host Miocene Monterey Group shales and cherts, and possibly deeper Cretaceous sediments of the Great Valley Group. MS4 water is more diluted by meteoric input than MS11. Long-term monitoring of these two springs shows they experience small seasonal fluctuations in temperature and that they are generally hydrologically and geochemically stable (**Fig. 1D-F**). Water temperatures of 27-29 °C are well above the mean annual surface temperature of 15.1 °C. The salinity of the springs is 1.8 and 2.3% for MS4 and MS11, respectively. The sulfide levels (within the zone of oxygenation) range up to ∼9 and 69 μmol/L at MS4 and MS11, respectively.

The biofilms at both MS4 and MS11 sites (**Fig. 1B-C**) are mainly composed of thin white streamers (∼ 5-10 cm long) that are primarily attached to rocks and Scanning electron microscopy (SEM) and scanning transmission X-ray microscopy (STXM) revealed that MS4 biofilms consist of filaments and cells distributed amongst the filaments (**Fig. 2**). By contrast, the MS11 biofilm consists almost entirely of filamentous bacteria (**Fig. 2C, Fig. 3C-D, Fig. 4**).

**Figure 2.**
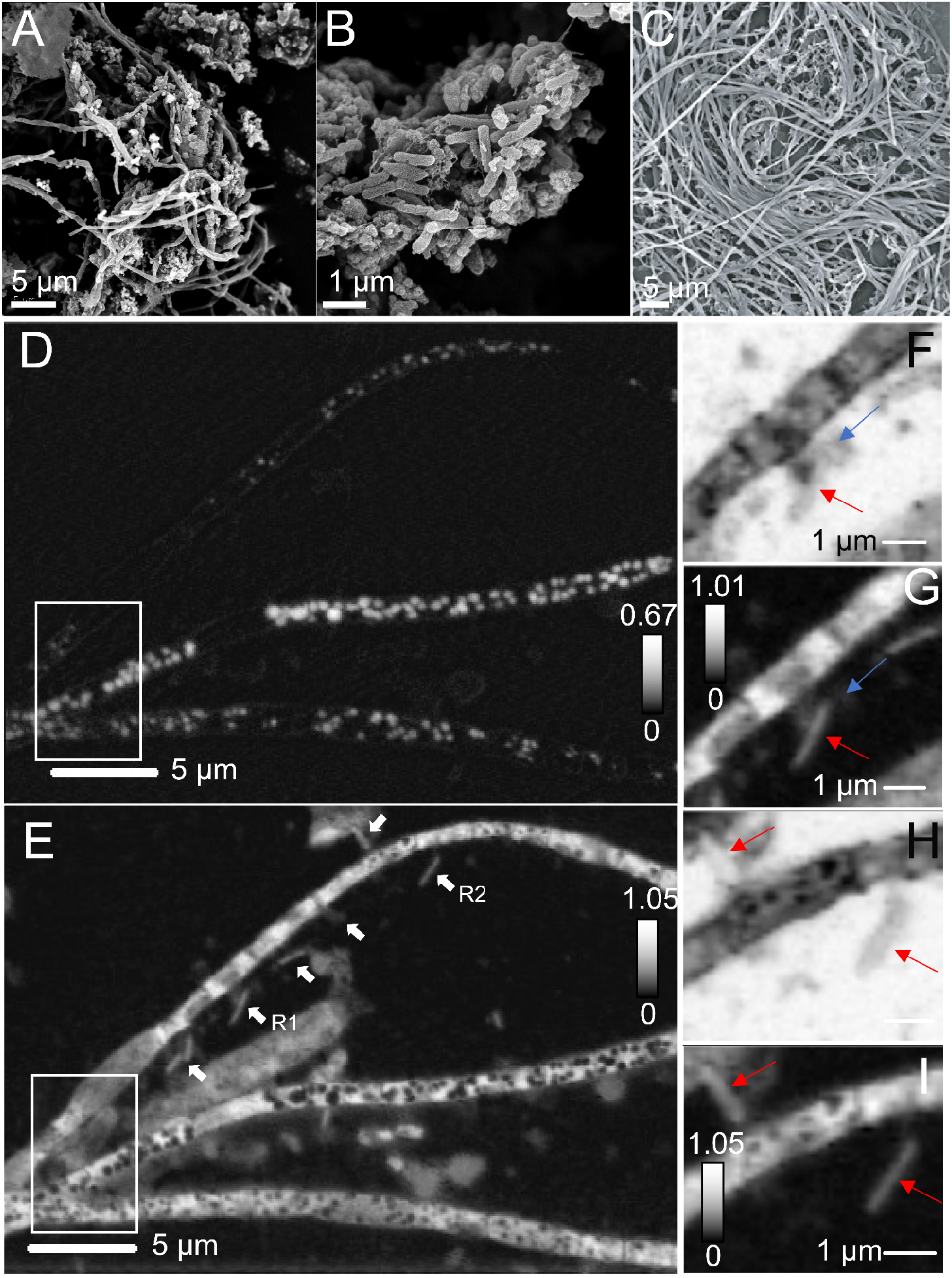
Microscopic characterization of the biofilms. A), B) Scanning electron micrographs of MS4 and C) MS11 biofilms. Scanning transmission x-ray microscopy of MS4 biofilms. D) Distribution map of S^0^ suggesting the presence of sulfur granules (378 ± 50 nm in diameter) within the compartments of the filaments. The width of top, middle and bottom filaments are 1.23 ± 0.48 µm, 1.01 ± 0.19 µm and 1.33 ± 0.3 µm respectively. E) Corresponding carbon map. White arrows point to cells. F) An ultra-small cell (476 ± 36 nm long, 246 ± 22 nm, blue arrow) in contact with an apparently episymbiotic cell (red arrow), imaged at 280 eV (region R1, panel E) and corresponding G) Carbon map. H) Two apparently episymbiotic cells (red arrows) connected to filaments, imaged at 280 eV (region R2, panel E) and corresponding I) carbon map. The intensity scales correspond to the optical density.

**Figure 3.**
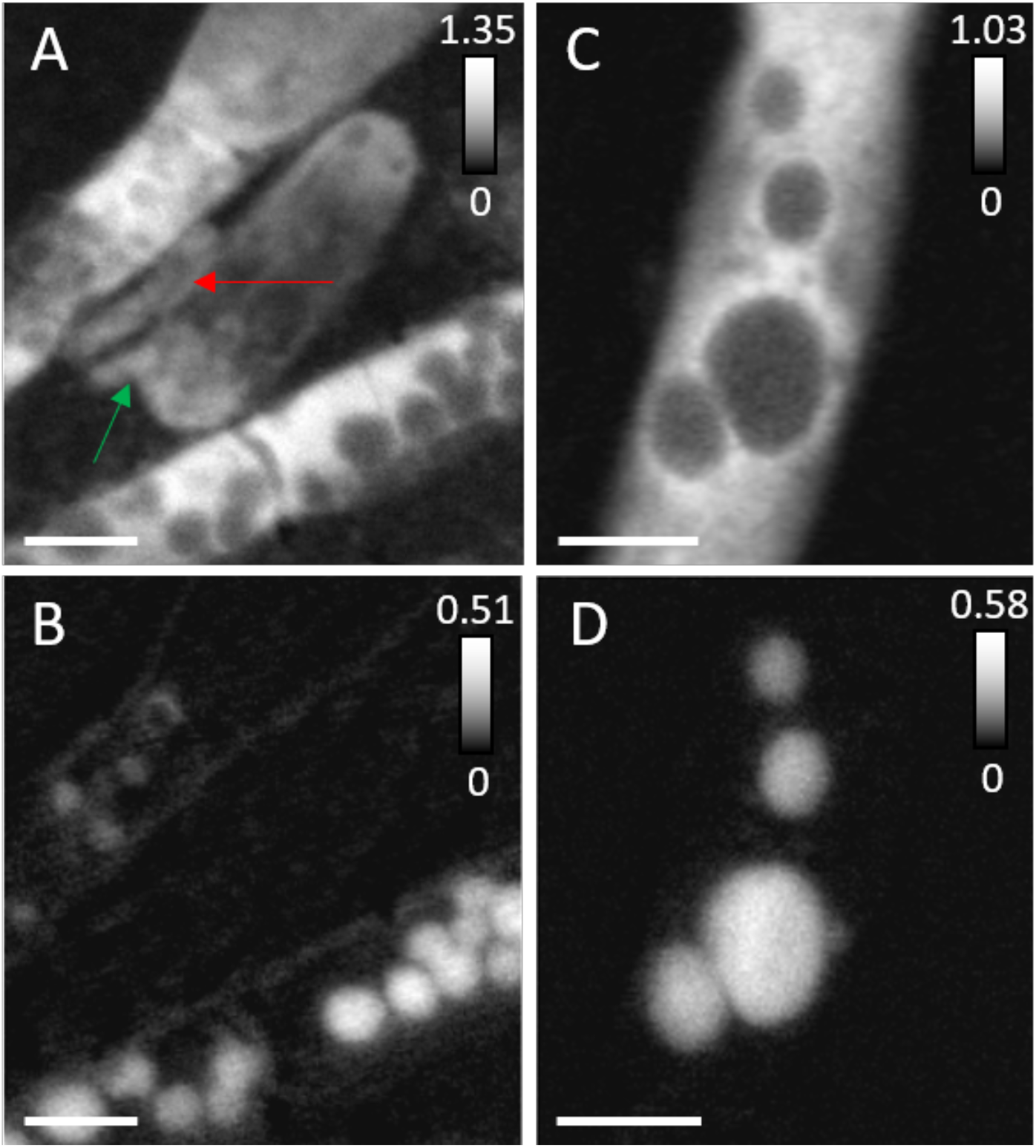
Scanning transmission x-ray microscopy of MS4 and MS11 biofilms. A) Protein map and corresponding B) Distribution map of S^0^ in MS4 biofilms (in white boxed area, Fig. 2). Cells that are 893 ± 29 nm long, 370 ± 20 nm wide (red arrow), 657 ± 30 nm long, 242 ± 32 nm wide (green arrow), seen in close contact with filaments. C) Protein map and corresponding D) Distribution map of S^0^ in MS11 biofilms, showing the presence of large sulfur granules (∼180 nm to ∼1.2 µm in diameter) in a small area of a long filament. The intensity scale corresponds to the optical density. Scale bars are 1 micron.

**Figure 4.**
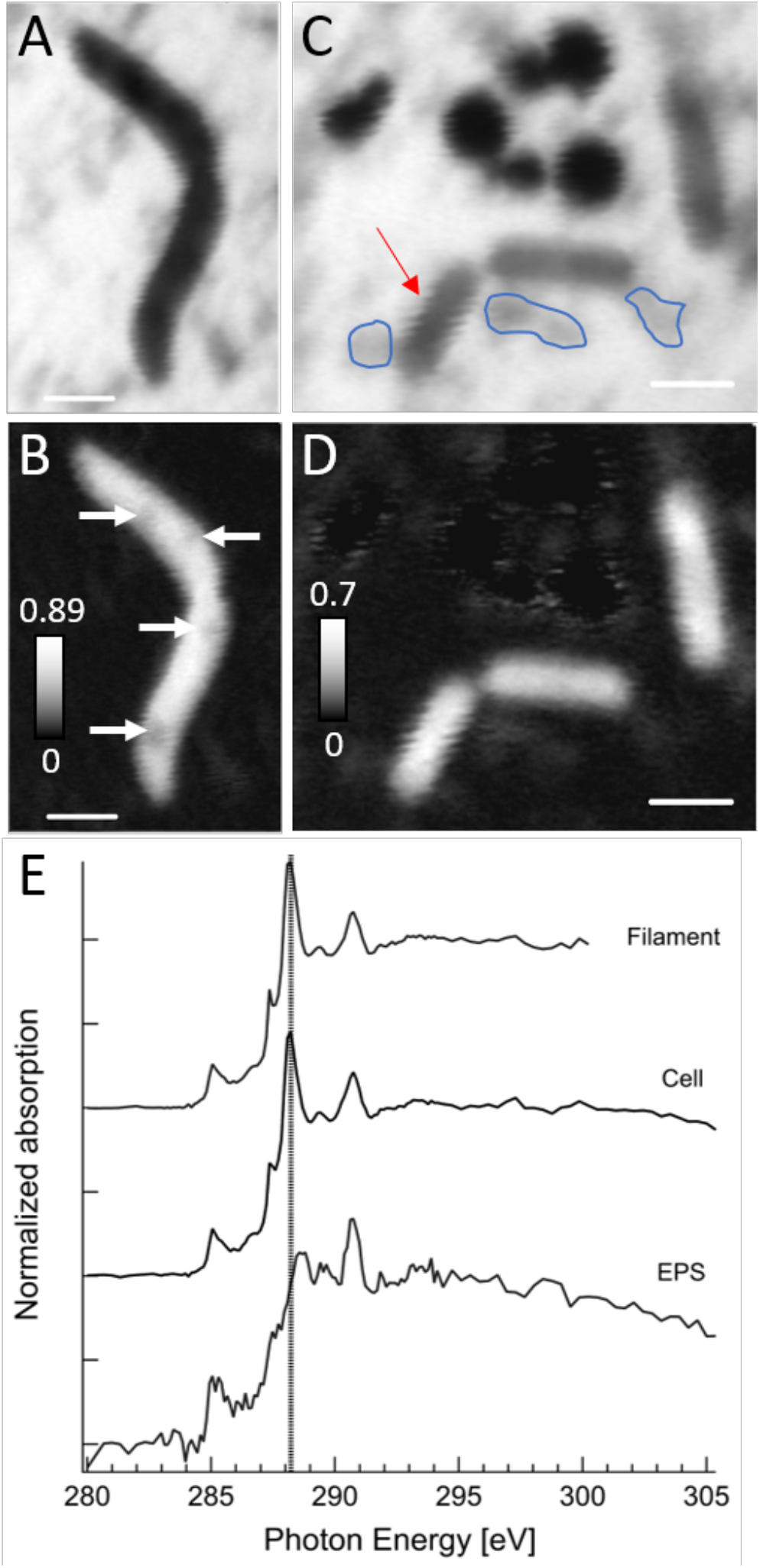
Scanning transmission x-ray microscopy at 87° Kelvin of frozen-hydrated MS11 biofilms. A) A small filament imaged at 288.2 eV (peak of the amide carbonyl groups in proteins) and corresponding B) protein map, granules are pointed by white arrows. C) Extracellular S^0^ granules (∼300 to 850 nm in diameter) near cells imaged at 288.2 eV and corresponding D) protein map. The intensity scale corresponds to the optical density. E) Carbon K-edge NEXAFS spectra of the filament (S^0^ granule-free areas), exhibiting a major peak at 288.2 eV, of a cell (red arrow) with main peak at 288.2 eV and of extracellular polymeric substances (EPS, circled in blue) exhibiting a main peak at 288.7 eV (carboxyl groups in acidic polysaccharides), see **Table S2** for details. Dashed line is at 288.2 eV. Scale bars are 1 micron.

### Filamentous bacteria have encapsulated elemental sulfur granules and episymbionts

Micro-focused XRF mapping of sulfur distribution at 95 °K evidenced the presence of sulfur across MS4 biofilm filaments (**Fig. S1**). STXM sulfur maps and S L_2,3_ NEXAFS spectra showed that these filamentous bacteria contain S^0^ granules (average 378 ± 50 nm diameter, as estimated on 76 granules) encapsulated in protein-rich compartments (**Fig. 2D-I**, **Fig. 3A-B**, **Fig. S2**). The width of these filaments is <1.6 µm suggesting the presence of *Thiotrix spp*. type I (ref). Rod-shaped, curved-shaped and coccoid cells were found near the filaments in MS4 biofilms **(Fig. 2, Fig. 3, Fig. S3**). C K-edge NEXAFS spectra at 87 °K of filamentous bacteria in MS11 (**Fig. 4**) exhibit a major peak at 288.2 eV corresponding to amide carbonyl groups evidencing that filaments are protein-rich (**Supplementary Table S2)**. Protein maps of these filaments (**Fig. 3C, Fig. 4B**) suggest that sulfur granules are surrounded by proteins. The spectrum of cells exhibits a major peak at 288.2 eV (amide bonds), a peak at 285.2 eV attributed mostly to aromatic groups in proteins and a peak at 289.5 eV attributed to nucleic acids, consistent with prior studies at room temperature [41,62–64], see **Supplementary Table S2** for details. Resonances are more defined, likely due to reduced Debye-Waller thermal disorder at low temperature. Cells, filaments and extracellular polymeric substances (EPS) exhibited a shifted carbonate peak at 290.7 eV that corresponds to either organic carbonates or carbonate minerals[65], and originates mainly from dissolved carbonates and carbonate precipitates present in the groundwater at circumneutral pH (**Supplementary Tables S1, S2**). Cells and filaments both contained potassium, but not the EPS. Strikingly, ultra-small cells were found along the surfaces of the filaments in both MS4 biofilms (**Fig. 2F-G, Fig. 3, Fig. S3D)** and MS11 biofilms (**Fig. S3A**), these cells are typically about 480 nm long, 250 nm wide, as estimated from STXM images. Other ultra-small cells (290 ±20 nm long, 120 ±10 nm wide) were also found in the vicinity of the filaments but not on their surfaces.

### Biofilms contain diverse bacteria and archaea and include CPR bacteria

We used genome-resolved metagenomics to investigate microbial consortia, metabolisms and microbial interactions that underpin the Alum Rock communities. In total, we recovered 212 non-redundant genomic bins from the MS4 and MS11 samples (57 from MS11 and 155 from the biofilm + planktonic samples from MS4). Of these, 38 were classified as near-complete (>95%, **Supplementary Table S3**). Taxonomic affiliations of all of the bacterial genomes were established based on concatenated ribosomal protein trees (**Fig. 5A**).

**Figure 5.**
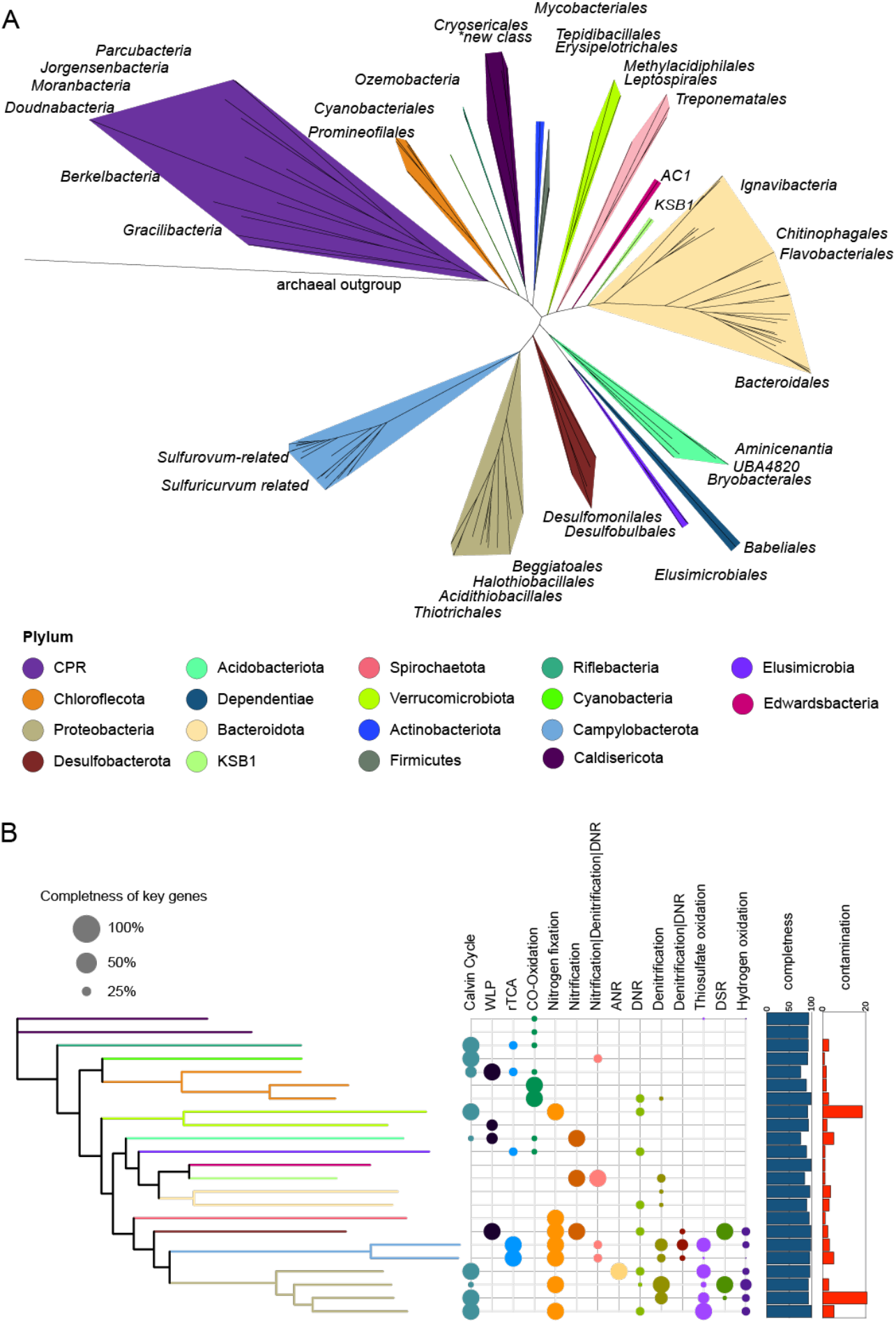
Phylogenetic analysis and metabolism of bacteria represented by MAGs from the MS4 and MS11 sites. A) The tree is based on 16 concatenated ribosomal proteins (rpL2, 3, 4, 5, 6, 14, 15, 16, 18, 22, 24 and rpS3, 8, 10, 17, 19) generated using iQ-TREE. An archaeon, *Thermoccocus alcaliphilu*s, was used as the outgroup. B) The metabolic capacities for generalized biogeochemical pathways in Alum Rock genomes are represented by colored circles. A pathway is present if the core KEGG orthologs encoding that pathway are identified in each genome. Abbreviations in the metabolic capaticites figure are as follows; WLP, Wood–Ljungdahl pathway, rTCA, eductive tricarboxylic acid cycle; ANR, Assimilatory nitrate reduction; DNRA, dissimilatory nitrate reduction to ammonia; Thiosulfate oxidation by SOX complex; DSR, Dissimilatory sulfate reduction; Hydrogen oxidation, [NiFe] hydrogenase and NAD-reducing hydrogenase.

Genomically represented groups in the biofilms and planktonic fractions from both sites include Gammaproteobacteria (Thiotrichales, Chromatiales, Beggiotales), Campylobacterota (Campylobacterales), Betaproteobacteria (including *Thiomonas*), Deltaproteobacteria (specifically Desulfobacterales), Bacteroidota, Chloroflexi, Ignavibacteria, Spirochaetes, Lentisphaerae, Riflebacteria, Verucomicrobia, Acidobacteria, Planctomycete, KSB1, Caldisericota, Planctomycetota, Edwardsbacteria, Dependentiae (TM6), and Margulisbacteria. Diverse groups of CPR are present, including Uhrbacteria (OP11), Gracilibacteria (BD1-5), Peregrinibacteria (PER), Moranbacteria (OD1), Woesebacteria (OP11), Roizmanbacteria and Gottesmanbacteria (OP11), Saccharibacteria (TM7), Falkowbacteria (OD1), Absconditabacteria (SR1), Berkelbacteria and Doudnabacteria and Dojkabacteria (WS6). (see: https://ggkbase.berkeley.edu/alumrock-genomes/organisms).

To estimate the abundances of organisms in the two springs (independent of binning) we calculated the DNA read coverage of ribosomal proteins from all of the genomic bins (**Fig. S4**). The MS4 spring was dominated by *Halothiobacillales, Beggiatoales and Thiotrichales* and, Campylobacterales based on relative abundance among genomes (**Supplementary table S4**). The most abundant species in MS4 shares genome-wide average 51% amino acid similarity with the sulfur oxidizer *Thiothrix nivea[66]*. The MS11 spring was dominated by a single *Beggiatoa* sp. (Beggiatoa-related_37_1401).

### Diverse bacteria are implicated in sulfur cycling

The MS4 biofilms are estimated to be ∼1.4 x as diverse as the MS11 biofilms, based on the number of phylogenetically informative marker genes detected (normalized for sequencing depth). We focused our analysis of the sulfur metabolism of MS4 bacteria for this reason, and given that we detected ultrasmall and surface-attached cells on filamentous bacteria implicated in sulfur oxidation. The most abundant organism in MS4, which is closely related to the filamentous bacterium *Thiothrix nivea*, encodes genes (*soxABC*, periplasmic thiosulfate-oxidizing; *aprAB*, adenylylsulfate reductase; *dsrAB*, reverse dissimilatory sulfite reductase) to convert sulfide to thiosulfate, elemental sulfur and sulfate (**Fig. 5B**). The absence of *dsrD* genes indicates that the Dsr complex operates in the sulfide oxidation direction (i.e. rDsr pathway). This *Thiothrix* bacterium also lacks any *soxC* genes, which in bacterial genomes has been associated with the accumulation of sulfur granules or polysulfide [67,68]. Based on the abundance of these organisms and their likely association with sulfur granules, it is possible that *Thiothrix* are the host for the ultra-small cells.

MS4 contains various other bacteria capable of oxidation of sulfur compounds. A subdominant population of *Sulfurovum* bacteria encode *sqr* genes and thus likely oxidize sulfide to S^0^. Some *Sulfurovum* bacteria in both communities have genomes also encode *soxCDYZ* complexes, suggesting they mediate thiosulfate oxidation (potentially coupled to nitrate reduction, e.g., via *narG* and *napA. Sulfuricurvum* species are also relatively abundant in MS4 and encode genes for sulfur and thiosulfate oxidation, in line with culture-based studies [69]. [69]. The genomes of *Chloroflexota* encode the capacity for thiosulfate disproportionation via thiosulfate reductase / polysulfide reductase (*phsA*) and sulfide oxidation via flavocytochrome *c* sulfide dehydrogenase. Two low abundance Gammaproteobacteria species related to *Acidthiobacillus* have the capacity for thiosulfate oxidation. Several genomes from moderately abundant *Halothiobacillales* have the metabolic capacity for sulfide and thiosulfate oxidation via *fccB, dsrAB* and *soxBCY* respectively (**Supplementary Table S5, S6**).

Some bacteria from MS4 spring also potentially mediate dissimilatory sulfate reduction. Specifically, the genomes of some *Desulfobacteriales* belonging to the families of *Desulfatiglandaceae, Syntrophobacterales, Desulfurivibrionaceae* and *Desulfarculales* encode the capacity to reduce sulfate back to sulfide via Dsr genes, likely coupled to oxidation of organic carbon or H_2_. Some rare Desulfocapsaceae from MS4 that are related to bacteria of the genus *Desulfocapsa* have thiosulfate reductase, group Group 3b [NiFe] (Hyd; possibly sulfhydrogenase), as well as SAT and APR for the oxidation of sulfite to sulfate. Thus, it appears these bacteria are involved in sulfur disproportionation whereby S^0^, thiosulfate, and sulfite are converted to H_2_ S and sulfate., as has been demonstrated in cultures of bacteria from this genus[70]. Other *Desulfocapsa* spp. have tetrathionate reductase genes, suggesting they are capable of converting tetrathionate to thiosulfate. The *Desulfocapsa*-related bacteria also contain *dsrABD* genes, which fall within the reductive cluster closely related to those from *Desulfocapsa sulfexigens*. We infer that the *Desulfocapsa*-related bacteria are capable of S disproportionation, as reported previously [71]. This presence of *dsrD* suggests that the species in the spring is capable of sulfate reduction. Only members of the candidate phylum Riflebacteria, family Ozemobacteraceae, have the capacity of anaerobic sulfate reduction via anaerobic sulfite reductase system (*asrABC*). A bacterium from a new class of *Caldithrix* from the MS4 spring is predicted to perform sulfur oxidation via dissimilatory sulfite reductase, sulfite oxidation, sulfate reduction and thiosulfate disproportionation (**Supplementary Table S5, S6**). We also identified abundant bacteria from novel families of Bacteroidetes, which generally encode thiosulfate reductase genes (*phS*) and adenylylsulfate reductase (*aprA*) involved in thiosulfate disproportionation and sulfate reduction.

Surprisingly, we identified persulfide dioxygenase (sdo) and rhodonase (thiosulfate sulfurtransferase) in genomes of *Elusimicrobia, Riflebacteria, Oscillatoriophycidae* and in a novel family of *Syntrophales* (**Fig. 6A**). These enzymes are also present in the mitochondria of plants and animals, as well as in a number of heterotrophic bacteria, where they play important roles in the detoxification of intracellular sulfide and sulfur assimilation respectively [72,73]. We also found a putative sulfur dioxygenase encoded in a Doudnabacteria genome that clusters with protein sequences of other CPR bacteria from public data. In the operon there is adjacent a sulfur transferase, suggesting its potential function in thiosulfate oxidation (**Fig. 10**). This is interesting because persulfide dioxygenase has not been linked to CPR bacteria previously. Modeling of the persulfide dioxygenase from Doudnabacterium using AlphaFold2 indicates that it has structural homology with the biochemically characterized persulfide dioxygenase (**Fig. 6B-D**). We identified these two adjacent genes in the genomes of several other CPR from high sulfide environments, including Kaiserbacteria (groundwater from California), Pacebacteria (wastewater), Moranbacteria, and Gracilibacteria (Crystal Geyser aquifer). Thus, we suggest that these genes may enable a variety of CPR bacteria to grow and generate energy from sulfur oxidation.

**Figure 6.**
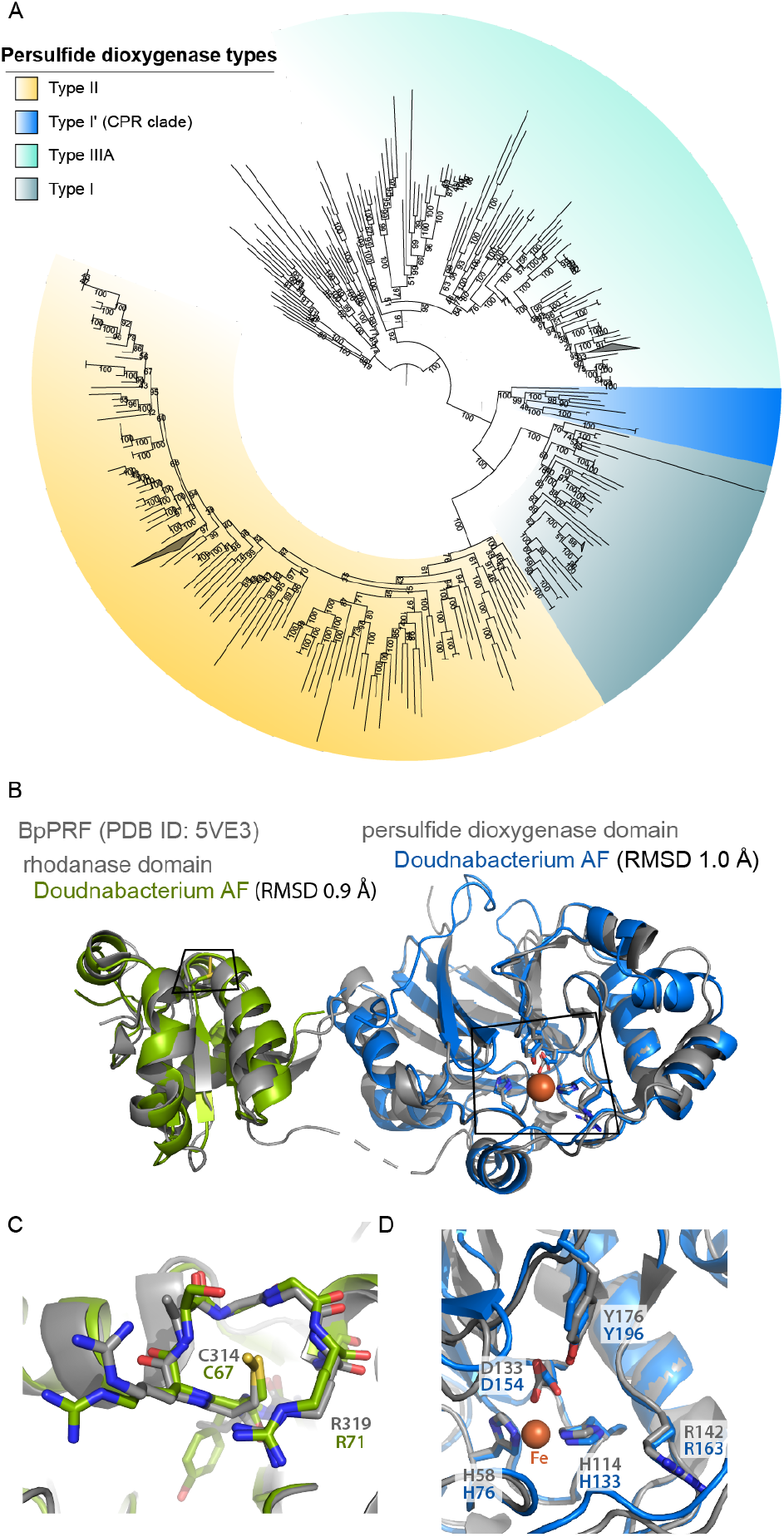
Novel Persulfide dioxygenase within CPR Bacteria. A) Phylogenetic analyses of persulfide dioxygenase proteins from the Alum Rock genomic bins. The blue monophyletic clade shows the persulfide dioxygenase found in CPR bacteria from sulfur-rich environments. B) AlphaFold models of Doudnabacterium putative rhodonase (green) and persulfide dioxygenase (blue) aligned with the corresponding domains of the characterized natural fusion protein BpRF (PDB ID: 5VE3). C) and D) Zoomed views of the active sites of the aligned structures reveal a strong coincidence of the key residues.

Like MS4, the most abundant microorganisms likely mediate sulfur compound oxidation, though*Beggiatoa* are the dominant species rather than *Thiothrix*. As expected, the *Beggiatoa* genome encodes a single contig that contains the Dsr genes (*dsrABPOJLCKMCHFE)*; s *dsrD* was not identified, we conclude that the Dsr genes are operational in a reverse Dsr pathway (rDsr). The genome also encodes AprAB (adenylylsulfate reductases), and Sat (sulfate adenylyltransferase) for the oxidation of sulfide to sulfate, sulfide-quinone oxidoreductase (Sqr) as well as sulfide dehydrogenase (*fccB*) genes for the oxidation of hydrogen sulfide to S^0^. The genomes do not contain a complete set of sulfur-oxidizing sox pathway genes, but *soxDXYZ* were identified. Given the lack of *soxC*, we conclude that (like *Thiothrix*) the primary role of *Beggiatoa* in the community is the conversion of sulfide to thiosulfate, elemental sulfur and sulfate. The absence of *soxCD* in bacterial genomes has been associated with the accumulation of sulfur granules or polysulfide [67,68].

### Sulfur oxidizing bacteria also contribute to nitrogen cycling

The dominant bacteria in MS4 and MS11 are predicted to mediate nitrogen fixation and denitrification processes. In both MS4 and MS11, genes encoding nitrogenase implicated in N_2_ fixation are widespread in Proteobacteria, including in the dominant *Thiothrix, Beggiatoa* and *Sulfurovum* and Verrucomicrobia. Other organisms with this capacity include other *Gammaproteobacteria, Chromatiales, Campylobacteriales, Sulfurovum, Sulfuricurvum, Ignavibacteria, Sulfosprillum, Spirochaetes, Desulfocapsa*, and potentially *Lentisphaerea*.

The *Thiotrichales* genomes encode numerous genes for the reduction of nitrate and nitrite, although the dominant *Thiothrix* species only has the capability to reduce nitrite to nitrous oxide via *nirS* and *norBC* genes. Some *Chromatiales* bacteria in both sites also appear to be capable of dissimilatory nitrite oxidation to ammonia. The sulfur-oxidizing Campylobacterales that occur in both MS4 and MS11 have numerous genes implicated in the reduction of nitrate (*napAB*) and nitric-oxide (*norBC*). Two low abundance *Acidithiobacillales* in MS4 that are predicted to perform thiosulfate oxidation have ammonia monooxygenase (*amoA*) genes, suggesting they may be involved in ammonia oxidation and nitrite ammonification. *Chloroflexi* that occur in both springs have the capacity for nitrite reduction via nitrite reductase (*nirK*), nitric oxide reduction (*norBC*) and nitrite ammonification. A novel *Caldithrix* species from MS4 has the potential of nitric oxide reduction via nitric oxide reductase (*norBC*) and nitrite reduction via periplasmic nitrate reductase NapA (**Fig. 5B**).

In addition to being the most abundant sulfur oxidizers in the MS11 spring, *Beggiatoa* are metabolically versatile with regards to nitrogen cycling. Their genomes encode genes with similarity to nitrate reductase (*narABG*), nitrite reductase (*nirS*), nitric oxide reductase (*norBC*), and nitrous-oxide reductase (*nosZ*) for the complete reduction of nitrate to N_2_. They also contain *nrfA* potentially for dissimilatory nitrite reduction to ammonia (DNRA) or nitrite ammonification. Thus, although these bacteria can grow aerobically, they also can likely couple sulfur oxidation to nitrate reduction, in line with previous studies.

### Extensive links between hydrogen and sulfur metabolism

To gain insight into the role of hydrogen metabolism in the Alum Rock springs, we analyzed the distribution of hydrogenases and associated enzymes in the genomes. There was considerable capacity for fermentative H_2_ production using nicotinamides (via group 3b and 3d [NiFe]-hydrogenases), ferredoxin (via group A [FeFe]-hydrogenases and group 4 [NiFe]-hydrogenases), and formate (via formate hydrogenases) as electron donors (**Fig. 7A**). Some putative H_2_ producers are likely to be metabolically flexible bacteria such as *Sulfurospirillum* and Flavobacteriales, which switch to fermentation when limited for respiratory electron acceptors based on previous reports [55,74], CPR bacteria, TA06, and Spirochaetes with group 3b and 3d [NiFe]-hydrogenases are likely to be obligate fermenters given they apparently lack terminal reductases (**Supplementary Table S7)**. The gene arrangements of the group 3b [NiFe]-hydrogenases in the genomes of the CPR bacteria *Berkelbacteria* and *Moranbacteria* (**Fig. 7B**) are similar to the biochemically characterized hydrogenase and sulfhydrogenase of *Pyrococcus furiosus* [75] and those previously reported in other CPR bacteria[25,76], suggesting that these hydrogenases may be capable of reversible oxidation of hydrogen or the reduction of sulfur compounds like polysulfide. We modeled the complex from *Berkelbacteria* genome using AlphaFold and the model suggests a hydrogenase module (α and γ subunits) with an electron wire of FeS clusters connecting to a nucleotide reducing module (β subunit) (**Fig. 7C**). The δ subunit has no close structural analogues but contains an additional FeS cluster and may accommodate an additional electron-accepting partner (**Fig. 7D**). Based on this structural analysis there are two separate paths for the electrons suggesting this 3b [NiFe]-hydrogenase complex is potentially an electron-bifurcating hydrogenase.

**Figure 7.**
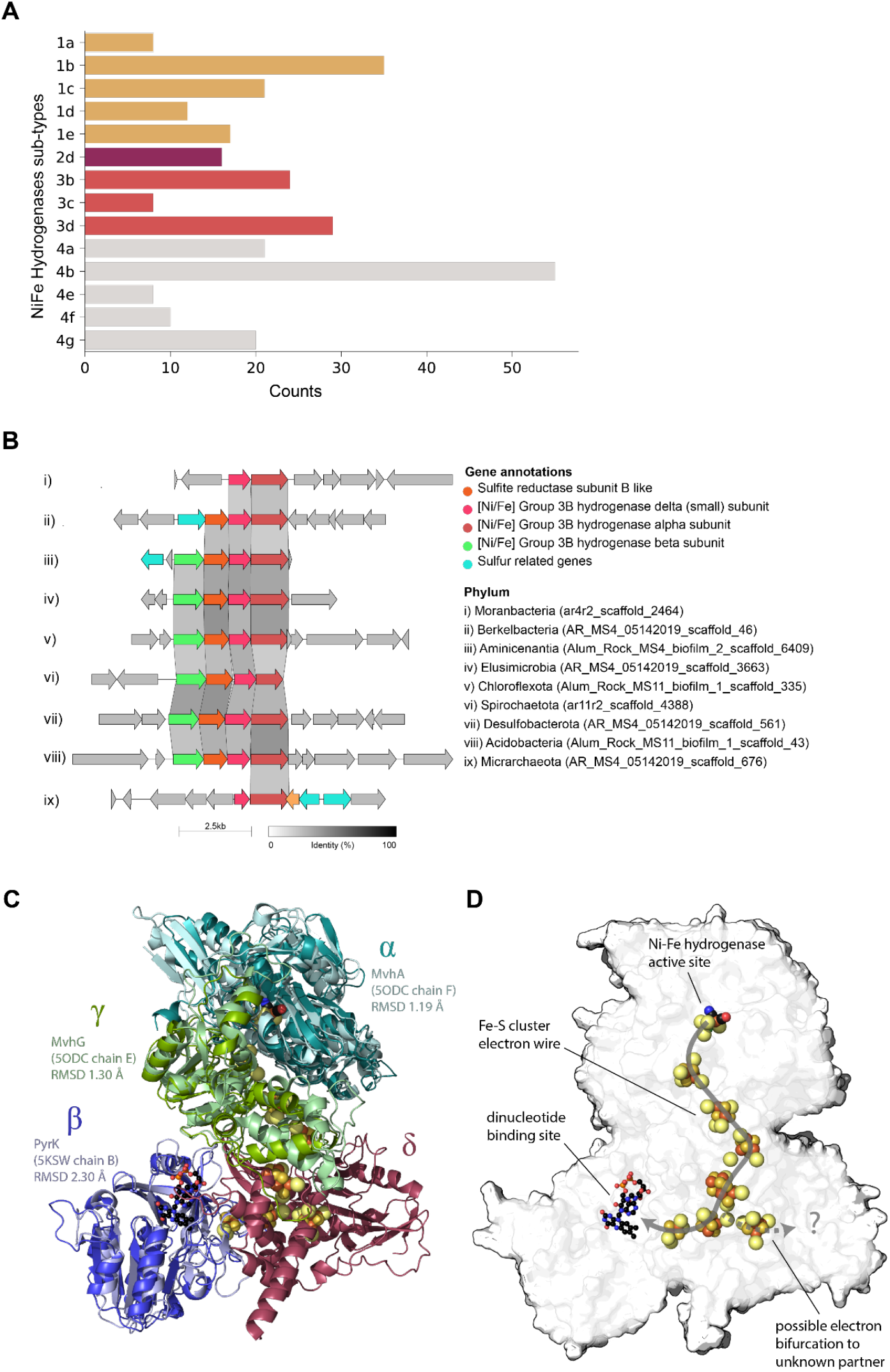
Hydroenases distribution in Alum Rock genomes and structural insights of Group 3b [NiFe]-hydrogenase complex. A) Total distribution of hydrogenases from the Alum Rock spring. B) Genomic organization of novel Group 3b [NiFe]-hydrogenases from different organisms present in the springs. C and D) Alphafold multimeric model for the Berkelbacterium putative Group 3b [NiFe]-hydrogenase complex with the closest known structural matches aligned to each protein.

Numerous bacteria in the Alum Rock springs are predicted to consume H_2_ for energy generation. Most of these hydrogenotrophs are predicted to use H_2_ to reduce sulfate (via group 1b and 1c [NiFe]-hydrogenases; primarily Deltaproteobacteria), elemental sulfur (via group 1e [NiFe]-hydrogenases; primarily Gammaproteobacteria), or heterodisulfides (via group 3c [NiFe]-hydrogenases; various lineages including Acidobacteria). The most abundant Gammaproteobacteria and Campylobacteria likely oxidize both H_2_ and sulfur compounds either mixotrophically or alternatively autotrophically. The hydrogenase repertoire of these organisms includes the oxygen-tolerant group 1b and 1d [NiFe]-hydrogenases [77,78].

### Organic carbon cycling and fermentation

The ability to fix inorganic carbon (CO_2_) is a common predicted capacity for bacteria from both sites (**Supplementary Table S5, S6**). The dominant *Thiothrix, Beggiatoa*, and *Chromatiales*-related bacteria have type II RuBisCO genes that function in the Calvin-Benson-Bassham (CBB) cycle (**Fig. S6)**. One Absconditabacteria genome has a RuBisCO that phylogenetic analysis places within the form II/III CPR clade, as reported previously [25,79]; these enzymes are inferred to function in a nucleoside salvage pathway in which CO_2_ is added to ribulose-1,5-bisphosphate to form 3-phosphoglycerate [80]. *Elusimicrobia* and *Campylobacterota*, including species related to Sulfurimonadaceae, have ATP citrate lyase genes that encode the key enzyme for CO_2_ fixation via the reverse TCA (rTCA) cycle. We also identified rTCA genes in a novel *Bacteroidetes* organism (**Supplementary Table S5, S6**). Genes of the Wood Ljungdahl carbon fixation pathway (*cooS/acsA, acsB and acsE*) were widespread in both springs, including in members of the Bacteroidetes, Desulfocapsa, Lentisphaerae, Chloroflexi, and Aminicenantia with the potential of oxidation of small organic compounds.

To infer polymer biomass degradation capacity of the biofilm organisms, we used marker genes involved in carbohydrate metabolism. Many bacteria in both springs have the capacity of hydrolyzing complex organic molecules to produce a varierity of electron donors such as acetate, hydrogen and lactate (**Fig. 8A**). Of the organisms in the community, *Bacteroidetes* and Ignavibacteria contain the most glycosyl-hydrolase genes and thus they likely play important roles in polysaccharide degradation. Notably, one *Bacteroidetes* from MS11 has 66 glycoside hydrolase genes. This organism is the only bacterium that appears to be capable of degrading cellulose, hemicellulose, polysaccharides, and monosaccharides. *Gammaproteobacteria, Spirochaetes, Bacilli, Lentisphaerae* also contain genes for the degradation of a variety of complex carbohydrates, but these genes are at relatively low abundance in the sulfur-oxidizing *Proteobacteria*. Similarly, many bacteria other than the sulfur-oxidizing *Proteobacteria* (and CPR) have indications of the capacity for beta-oxidation pathway of saturated fatty acids to acetyl-CoA. Many of the CPR bacteria have a few glycosyl hydrolase genes, which is significant given the scarce indications of other metabolic capacities in these organisms. Methane oxidation is predicted to be a capacity of members of Verrucomicrobia, specifically members of the Methylacidiphilales. This reaction involves particulate methane monooxygenase (pMMO-ABC), the genes for which were identified and classified phylogenetically.

**Figure 8.**
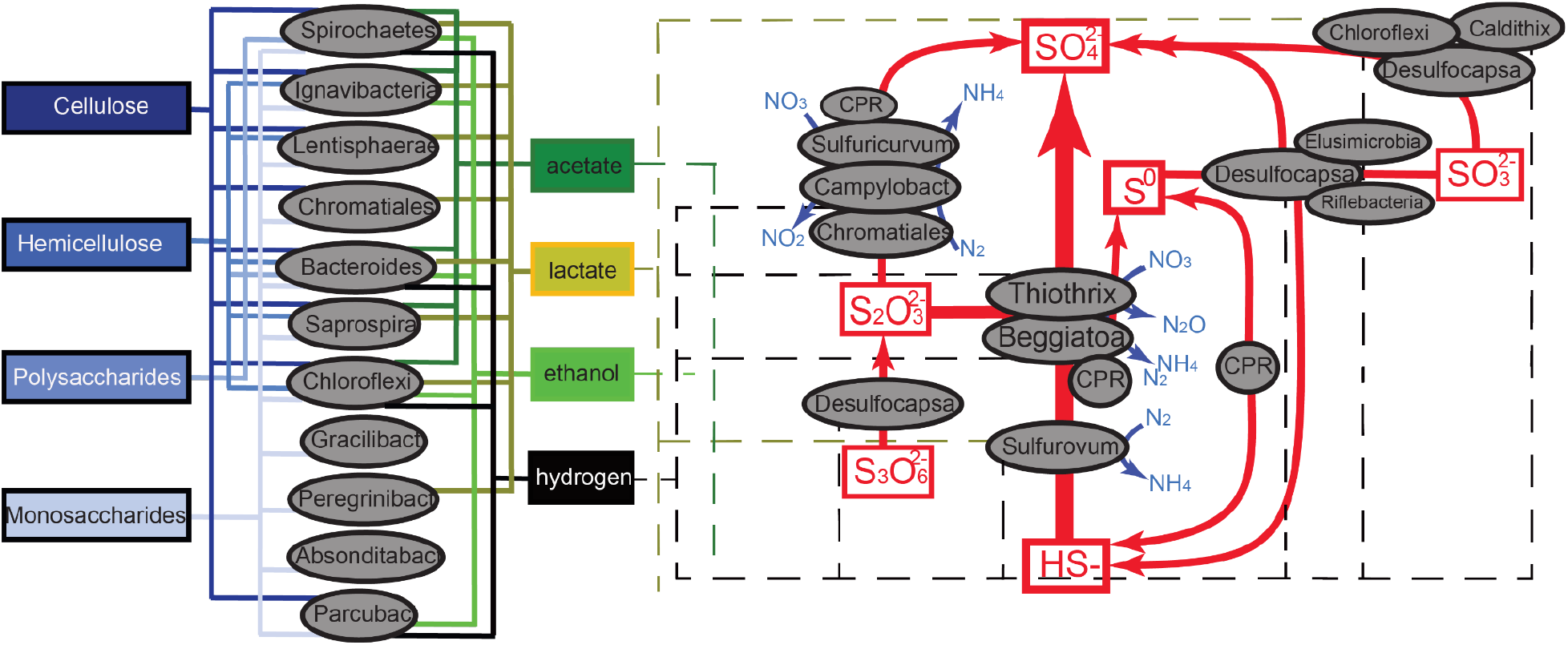
Inference of partitioning of carbon, sulfur and nitrogen cycling in the Alum Rock springs. Based on the gene content of genomes reconstructed from the springs. Arrows indicate metabolic capacities reconstructed from metagenomes recovered from the Alum Rock mineral springs. The dashed lines represent potential electron donors for anaerobic respiration processes.

One of the more interesting organisms present in the MS4 spring is a Gracilibacteria, which is predicted to have minimal metabolic capacities beyond glycolysis, production of peptidoglycan and generation of formate, some of which may be exported for use by other community members. Other capacities predicted for this bacterium are production of riboflavin, amino-sugars, RNA degradation, 1C by folate, interconversion of purines and pyrimidines and biosynthesis of a few amino acids.

### Phages may contribute auxiliary metabolic genes

We genomically sampled 36 dsDNAv phages (**Supplementary Figigure S7**) (28 from MS4 and 8 from MS11) to and one nucleocytoplasmic large DNA phage. These phages have genes potentially involved in translation (bacterial ribosome L7/L12 and ribosomal protein S1), nitrogen utilization, carbon metabolism, iron metabolism (ferritin), and nucleotide metabolism (pyrimidine deoxyribonucleotide and adenine ribonucleotide biosynthesis) and defense systems such as CRISPR-Cas and TROVE (Telomerase, Ro and Vault module).

## Discussion

Some springs are hotspots where resources associated with deeply sourced water can sustain chemoautotrophic ecosystems independent of sunlight. We studied two closely spaced but distinct sites that discharge a mixture of deeply sourced and shallow groundwater, providing microorganisms with both reduced compounds and access to oxygen. Our research integrated geochemical, X-ray spectromicroscopy, and genome-resolved metagenomic data to resolve the network of microorganisms that define the ecosystems. This approach provided insights into organism associations, including those that involve CPR bacteria, and the biogeochemical processes that sustain autotrophic ecosystems in the context of their spring-based hydrological setting.

Analysis of the metabolisms of the dominant bacteria in the springs revealed that genes implicated in sulfur cycling are common at both sites (**Fig. 8B**). As expected, the main energy source is reduced sulfur in the form of sulfide. Overall, the most common sulfur metabolisms are sulfide oxidation, thiosulfate disproportionation, sulfur oxidation, and less commonly sulfite oxidation and sulfate reduction. Sulfide can be oxidized aerobically and in some cases anaerobically, coupled with nitrate reduction. The genomic analyses suggest that intermediate sulfur compounds, as well as sulfate and sulfide, are actively cycled by Campylobacterota (*Sulfurovum, Thiovulum*), Gammaproteobacteria (*Thiotrichales and Beggiotales*) in the spring communities, probably coupled to nitrogen compound reduction in some microhabitats. Partly oxidized sulfur in the form of elemental sulfur likely serves as an energy source that is stored as sulfur granules. Interestingly, elemental sulfur-bearing granules within filamentous cell compartments of *Beggiatoa* and/or *Thiothrix* likely serve as an energy source for the growth of these bacteria. The sulfur oxidizers are the primary source of fixed carbon and nitrogen.

A higher flow rate and a higher concentration of sulfate was observed at MS11 compared to MS4, and the communities have distinct microbial characteristics (**Supplementary Figure S3**). The MS4 ecosystem is highly diverse and dominated by abundant sulfide-oxidizing Gammaproteobacteria (*Thiothrix, Sulfurovum*) and sulfate-reducing Desulfobacteriales. The MS11 spring has relatively low diversity and is highly dominated by Campylobacterota (*Sulfurovum, Thiovulum*) and Gammaproteobacteria (*Thiotrichales and Beggiotales*). Our findings are consistent with predictions from studies that indicate that filamentous Campylobacterota dominate biofilms with high sulfide/oxygen (>150) ratios whereas Gammaproteobacteria (*Beggiatoa*-like) prefer lower (<75) ratios[9].

We focused some analyses on the diverse CPR bacteria within these communities, as their roles in sulfur-based chemoautotrophic ecosystems remain poorly known. CPR bacteria are characterized by small genomes and minimal anaerobic fermentative metabolism[81], however recent studies have shown auxiliary metabolisms such as the presence of hydrogenases[25,76], rhodopsin[82], nitrite reductases[83] and F-type ATPase[84], that may contribute to alternative energy conservation and adaptations to different environments and host associations. Notably, we identified genes potentially involved in elemental sulfur reduction (Sulfyhydrogenase) and thiosulfate oxidation (persulfide dioxygenase and rhodonase) in genomes of some CPR bacteria, suggesting a potential new energy generation mechanism for these bacteria. We found that other CPR from high sulfur environments have the same predicted potential for thiosulfate oxidation, suggesting an important general adaptation of CPR bacteria in sulfur-rich environments.

Perhaps the most interesting aspect of the current study regards interactions involving CPR bacteria and their host microorganisms. CPR-host associations have rarely been documented, with the exception of oral microbiome-associated Saccharibacteria (TM7) [29,85] and Actinobacteria. For this association, laboratory studies[86] have validated genomic predictions of metabolic interdependency[76]. One study imaged the CPR cells on the surfaces of their Actinobacteria hosts via SEM and showed them to be rod-shaped and < 0.2 µm in diameter and ∼0.5 µm in length[87]. Another study linked *Vampirococcus* with anoxygenic photosynthetic Gammaproteobacteria[88]. Two studies suggest links between Parcubacteria and archaea, in one case *Methanosaeta[89]* and *Methanothrix[89]*. In the case of the Nealsonbacteria CPR associated with *Methanosaeta*, cryo-TEM images indicate that the cells are ∼0.5 µm in diameter. Other cultivation-independent studies have verified that CPR cells are ultra-small, so can be enriched via filtration through a 0.2 µm filter[81]. Cryo-TEM images and tomographic analyses have documented ultra-small cells in direct association of CPR cells and host bacteria[31,81]. Generally, these data indicate that CPR cells are a fraction of a micron in length and diameter, consistent with the size for filament-associated ultra-small cells reported here (∼600 nm long, ∼200 nm width). Thus, we conclude that the ultra-small cells imaged in the MS4 biofilm are CPR bacteria.

Here, STXM imaging and NEXAFS spectroscopy of MS4 biofilms revealed the putative CPR bacterial cells occur in close proximity to filamentous cells with large sulfur granules. We infer that these filamentous cells are probably *Thiothrix*, given that they appear to be the only abundant filamentous bacteria in this sample and that they have the genomic capacity for sulfur-oxidation, including the capacity to produce elemental sulfur. Given the combination of imaging and genomic information, we predict that certain CPR cells are episymbionts of filamentous sulfur-oxidizing *Thiothrix*. Likely CPR identifications include Gracilibacteria, Berkelbacteria, Moranbacteria or Doudnabacteria, based on microbial community abundance information. Co-cultivation of *Thiothrix* and their episymbionts is needed to identify the CPR types, and to better understand the nature of their association (*e*.*g*., mutualistic, parasitic). Although only based on *in vitro* data from *Pyrococcus[75,90]*, the prediction that some CPR bacteria have the capacity to produce H_2_ S raises the possibility that these episymbionts are involved in cryptic sulfur cycling that involves sulfur-oxidizing bacteria. If so, it seems plausible that Berkelbacteria or Moranbacteria, which may be able to produce H_2_ S, are the CPR episymbionts that were imaged in this study.

Hydrogen is an important resource in many environments[91], yet little is known about the distribution and importance of hydrogenases in sustaining groundwater microbiomes. The most common chemolithoautotrophs in the Alum Rock spring biofilms are H_2_ -oxidizing bacteria, which use H_2_ as an energy source via the enzyme hydrogenase. Specifically, group 3b [NiFe]-hydrogenases are widely distributed in the genomes of many of the microbial community members. These complexes may mediate hydrogen metabolism or the direct hydrogenation of elemental sulfur to hydrogen sulfide [90]. Other hydrogenases of the microbial community members are implicated in hydrogen production and oxidation. Together, these findings suggest that most bacteria in Alum Rock springs cycle hydrogen gas and sulfur compounds, reactions that underpin the biology and geochemistry of this ecosystem.

## ADDITIONAL INFORMATION AND DECLARATIONS

### Competing Interests

JFB is a co-founder of Metagenomi.

#### Author contributions

**L.E.V-A** was involved in metagenome sample preparation, genomic and metabolic reconstruction, phylogenetic and protein structure analyses, data integration and writing of the paper. **S.C.F** was involved in STXM and X-ray fluorescence microprobe sample preparation and data analysis, data integration and writing the paper. **C.G**. contributed to the hydrogenases analyses and writing the paper. **A. J. P**. contributed to analyses of the genomic data. **A.L.J**. contributed to analyses of the CPR metabolism. **J.W-R**. contributed to phylogenetic analyses. **M.M**. helped conceive the study, collected water samples and measurements, and analyzed the geochemical data. **J.R**. collected the geochemical data. **L.M.O**. contributed to protein structure modeling and analyses **B.J.B**. contributed to phylogenetic and metabolic reconstruction and analyses. **D.F.S**. contributed financial support. **J.F.B**. conceived the study, was involved in writing the paper and analyses of the genomic data.

### Code and data availability

The Alum Rock genomes and raw sequencing reads for this study will be made available under NCBI BioProject number XXXX. The genomes presented in this manuscript are also made available at https://ggkbase.berkeley.edu/alumrock-genomes

## Acknowledgments

We thank Christine He, Raphaël Méheust, Susan Mullen, Ben Rubin, Jordan Hoff, Lily Law and Haridha Shivram for their assistance with one sampling trip. Rohan Sachdeva, Shufei Lei, Alex Crits-Christoph, Adair Borges for helpful discussions, and comments on the manuscript. We thank Tolek Tyliszczak for help with the STXM cryostage. We thank Alum Rock Park for granting sampling permits.

